# Impact of rapid susceptibility testing and antibiotic selection strategy on the emergence and spread of antibiotic resistance in gonorrhea

**DOI:** 10.1101/122200

**Authors:** Ashleigh R Tuite, Thomas L Gift, Harrell W Chesson, Katherine Hsu, Joshua A Salomon, Yonatan H Grad

## Abstract

**Background:** Increasing antibiotic resistance limits treatment options for gonorrhea. We examined the extent to which a hypothetical point-of-care (POC) test reporting antibiotic susceptibility profiles could slow the spread of resistance.

**Methods:** We developed a deterministic compartmental model describing gonorrhea transmission in a single-sex population with three antibiotics available to treat infections. Probabilities of resistance emergence on treatment and fitness costs associated with resistance were based on characteristics of ciprofloxacin, azithromycin, and ceftriaxone. We evaluated time to 1% and 5% prevalence of resistant strains among all isolates with: (1) empiric treatment (azithromycin plus ceftriaxone), and treatment guided by POC tests determining susceptibility to (2) ciprofloxacin only and (3) all three antibiotics.

**Findings:** Based on current gonococcal susceptibility patterns in the United States, the model indicated that continued empiric dual antibiotic treatment without POC testing resulted in >5% of isolates being resistant to both azithromycin and ceftriaxone within 15 years. When either POC test was used in 10% of identified cases, this was delayed by 5 years. The three antibiotic POC test delayed the time to reach 1% prevalence of triply-resistant strains by 6 years, while the ciprofloxacin-only test resulted in no delay. Results were less sensitive to assumptions about fitness costs and test characteristics with increasing test uptake. The main limitation of this study is that we made simplifying assumptions to describe gonorrhea transmission and the emergence and spread of resistance in the population.

**Conclusions:** Rapid diagnostics that report antibiotic susceptibility have the potential to extend the usefulness of existing antibiotics for treatment of gonorrhea. Monitoring resistance patterns will be critical with the introduction of such tests.

## INTRODUCTION

Increasing antibiotic resistance poses an immense challenge to the clinical and public health community [1], and underscores the importance of developing new strategies to control resistance. Among the most urgent threats to our ability to treat infections is antibiotic resistant gonorrhea. Treatment of gonorrhea is almost always empiric, because diagnosis is most commonly made by nucleic acid amplification test, which provides no susceptibility data. Even when culture is available, clinical and public health principles demand rapid treatment of patients without waiting for antibiotic susceptibility results.

Gonorrhea treatment guidelines are based on population resistance surveys, with antibiotics no longer recommended once resistance prevalence exceeds 5%. Only ceftriaxone and azithromycin remain as first-line therapy [2], and increasing resistance to both has been observed [1]. With empiric treatment strategies, a majority of gonococcal infections may remain susceptible to antibiotics no longer recommended for use. For example, 81% of gonococcal infections in the United States are susceptible to fluoroquinolones [3].

Given the prevalence of susceptible isolates, one proposed strategy to control resistance is the use of rapid diagnostics that allow clinicians to tailor treatment to the antibiotic susceptibilities of individual infections [4–7]. Sequence-based diagnostics for fluoroquinolone susceptibility report high positive and negative predictive values [8, 9] and are moving toward use in clinical care [10]. Large-scale genome sequencing is generating data on the test characteristics of nucleic acid-based diagnostics for susceptibility to other antibiotics [11, 12].

Underlying the promise of rapidly determining antibiotic susceptibility is the hypothesis that tailored therapy will prolong the utility of anti-gonococcal agents and better control resistance than empiric treatment. Here, we used a mathematical model of gonorrhea transmission to test the impact of tailored therapy on the prevalence of gonococcal infection and antibiotic resistance to three commonly used drugs: ciprofloxacin, azithromycin, and ceftriaxone.

## METHODS

### Model overview

We developed a dynamic compartmental model that describes gonorrhea transmission in a single sex population stratified by sexual risk. This model represented a population of men who have sex with men (MSM), who experience a significant burden of gonorrhea in the United States and in whom emergence of resistance is of concern [3, 13]. The natural history of gonorrhea infection was described by the following states: susceptible, symptomatic infection, and asymptomatic infection (Figure 1). Each of the infectious states was further subdivided to represent the resistance profile of the infecting strain. Model parameters are presented in Table 1 and additional model details are provided in the Technical Appendix.

**Figure 1.**
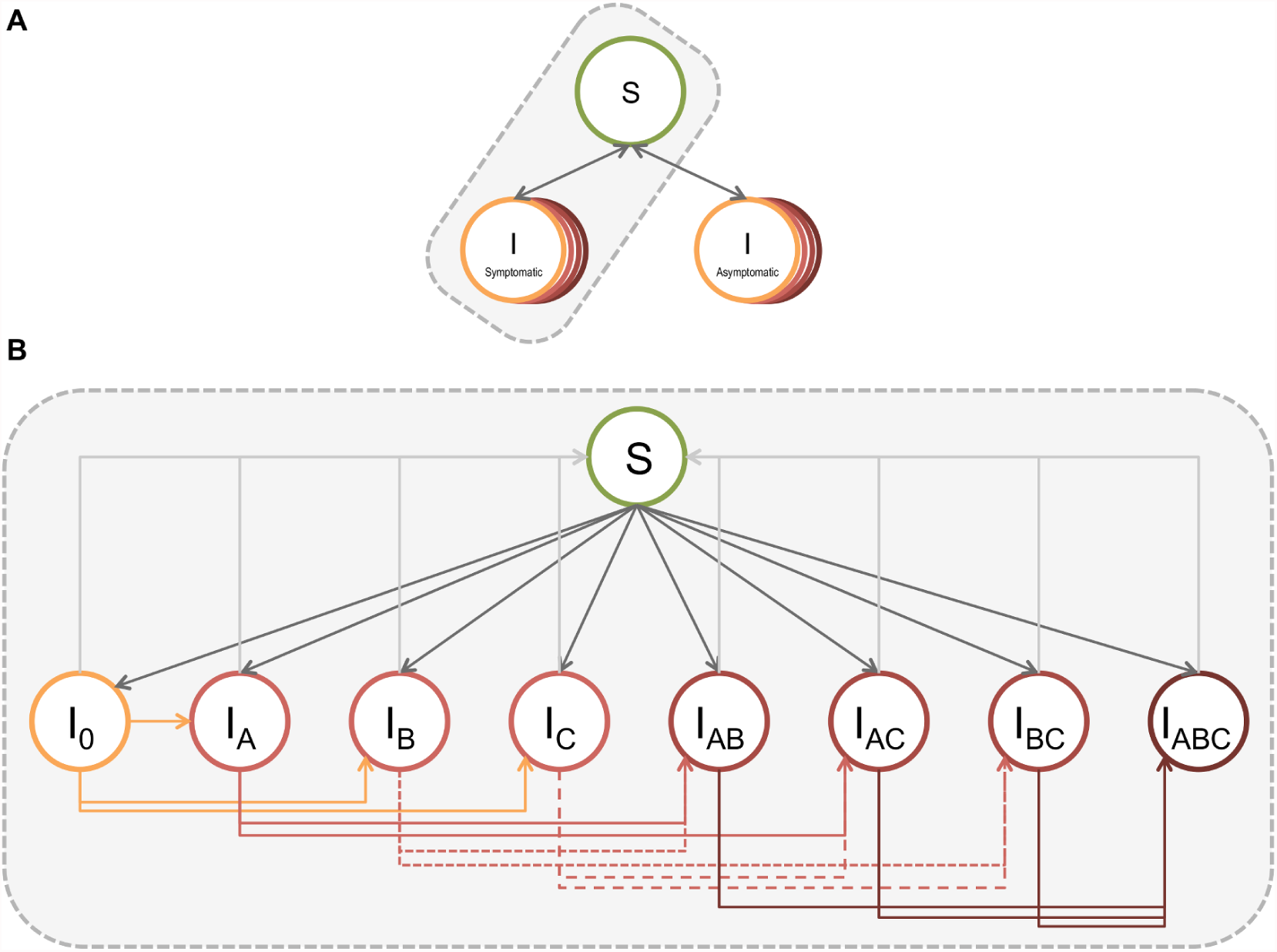
Overview of gonorrhea transmission model. (A) The model includes three states: susceptible, symptomatic infectious, and asymptomatic infectious. Infected individuals can return to the susceptible state via treatment or natural clearance of infection. (B) Expanded view of the different possible infected states, where subscripts indicate resistance to antibiotics A, B, and/or C. *I*_0_ indicates infection with a completely drug susceptible strain. Note that the same series of infectious states and transitions exist for symptomatic and asymptomatic infections. The model is further stratified by three sexual activity classes.

**Table 1.** Model population, gonorrhea natural history, and treatment parameters.

| Parameter | Details | Symbol | Value | Source |
| --- | --- | --- | --- | --- |
| Population size | | N | $10^6$ | Assumption |
| Gonorrhea prevalence at start (%) | Calibration target |  | 2.3 (1.2-2.8) | [22–24] |
| Proportion of cases resistant to drug X at start of evaluation period | | $\theta$ | | [3] |
| | A (ciprofloxacin) | $\theta_A$ | 0.189 | |
| | B (azithromycin) | $\theta_B$ | 0.023 | |
| | C (ceftriaxone) | $\theta_C$ | 0.0001 | |
| | A and B | $\theta_{AB}$ | 0.0022 | |
| | A and C | $\theta_{AC}$ | 0.0009 | |
| | B and C | $\theta_{BC}$ | $10^{-5}$ | Assumption |
| | A, B, and C | $\theta_{ABC}$ | $10^{-6}$ | Assumption |
| Sexual risk group distribution |  | n |  | Assumption |
|  | High |  | 0.1 |  |
|  | Intermediate |  | 0.6 |  |
|  | Low |  | 0.3 |  |
| Relative rate of partner change in risk groups |  | rp |  | [25] |
|  | High |  | 20 |  |
|  | Intermediate |  | 5 |  |
|  | Low |  | 1 |  |
| Rate of partner change in low risk group (/y) | | $c_{min}$ | 1.16 | Model fitting |
| Mixing parameters | | $\epsilon$ | 0.23 | Model fitting |
| Rate of model entry/exit (y) | | $\rho$ | 1/20 | Assumption |
| Transmission probability per partnership | | $\beta$ | 0.44 | [26–32]; model fitting |
| Probability symptomatic infection | | $\sigma$ | 0.6 | Assumption and model fitting |
| Average duration of infection without treatment (y) | | $1/\delta$ | 0.5 | [28, 31–33]; model fitting |
| Average time to treatment (y) | Symptomatic | $1/\tau_s$ | 0.026 | [34]; model fitting |
| Screening rate (/y) | Asymptomatic | $1/\tau_m$ | 0.39 | Assumption and model fitting |
| Probability of retreatment with effective drug, if initial treatment failure |  |  |  | Assumption |
| | Symptomatic infection | $\pi_s$ | 0.9 | |
| | Asymptomatic infection | $\pi_m$ | 0 | |
| Treatment rate if initial treatment failure (/y) |  |  |  | Assumption |
| | Symptomatic infection | $\tau_{sr}$ | $\tau_s/3$ | |
| | Asymptomatic infection | $\tau_{mr}$ | $\tau_m/3$ | |

### Treatment

We modeled treatment with three different antibiotics, which could be used individually or in combination. Each antibiotic had a probability of resistance emergence on treatment and a fitness cost associated with resistance, where fitness refers to the capability of the pathogen to survive [14]. We modeled fitness cost in terms of transmissibility relative to the susceptible strain[15]. The properties of each of the antibiotics were selected to mirror the classes of antibiotics used to treat gonorrhea infection: fluoroquinolones (A), macrolides (B), and extended spectrum cephalosporins (C) [16] (Table 2). Details of the characterization of these properties are provided in the Technical Appendix. In the absence of a point-of-care (POC) test to determine strain susceptibility, antibiotic choice reflected U.S. treatment guidelines of combination therapy with azithromycin and ceftriaxone (B and C in our model) [2].

**Table 2.** Characteristics associated with point-of-care test and drug-resistant *N. gonorrhoeae* strains.

| Parameter | Details | Symbol | Value<br>(base case and range) |
| --- | --- | --- | --- |
| Probability of resistance with treatment | Antibiotic A | $\omega_A$ | $10^{-6}$ ($10^{-9} - 10^{-3}$) |
| | Antibiotic B | $\omega_B$ | $5 \times 10^{-7}$ ($10^{-9} - 10^{-3}$) |
| | Antibiotic C | $\omega_C$ | $10^{-8}$ ($10^{-9} - 10^{-3}$) |
| Relative fitness of resistant strain | Susceptible | $f_0$ | 1 |
| | Antibiotic A resistant | $f_A$ | 1 (0.85-1) |
| | Antibiotic B resistant | $f_B$ | 0.94 (0.85-1) |
| | Antibiotic C resistant | $f_C$ | 0.98 (0.85-1) |
| | Antibiotics AB resistant | $f_{AB}$ | 0.94 (0.72-1) |
| | Antibiotics AC resistant | $f_{AC}$ | 0.98 (0.72-1) |
| | Antibiotics BC resistant | $f_{BC}$ | 0.92 (0.72-1) |
| | Antibiotics ABC resistant | $f_{ABC}$ | 0.92 (0.61-1) |
| Point-of-care test sensitivity | Antibiotic A | $\kappa_A$ | 1 (0.5-1) |
| | Antibiotic B | $\kappa_A$ | 1 (0.5-1) |
| | Antibiotic C | $\kappa_A$ | 1 (0.5-1) |
| Point-of-care test specificity | Antibiotic A | $\psi_A$ | 1 (0.5-1) |
| | Antibiotic B | $\psi_A$ | 1 (0.5-1) |
| | Antibiotic C | $\psi_A$ | 1 (0.5-1) |

### Deployment of a rapid diagnostic to determine susceptibility

We compared empirical treatment to treatment guided by a hypothetical POC test that rapidly determines susceptibility to: (i) a single antibiotic (A) and (ii) all three antibiotics. Based on a case’s resistance profile, antibiotic treatment was selected (Table S1), with the probability that the chosen antibiotic effectively treated the infection dependent on the test characteristics.

*Scenario I*. POC test for determining resistance to antibiotic A only. Antibiotic A was used to treat A susceptible infections, while A resistant infections were treated with combination BC therapy.

*Scenario II*. POC test for determining resistance to antibiotics A, B, and C. If multiple antibiotics would be effective at treating a case (i.e., infected with a completely susceptible strain or a strain resistant to only one antibiotic), we treated with the antibiotic with the highest fitness cost associated with resistance.

In scenario II, if an individual was identified as having a triply resistant infection, we assumed that the infection was ultimately successfully treated, with an alternative agent or higher antibiotic doses [17–19].

We evaluated different levels of POC test uptake in the population. When susceptibility was not determined prior to treatment, we assumed treatment according to guidelines, as described above.

### Model fitting

We calibrated model parameters describing gonorrhea natural history and sexual behavior using maximum likelihood estimation. Details are provided in the Technical Appendix.

### Model outputs and analysis

The model was initiated at the equilibrium prevalence determined through model fitting in the absence of resistant strains. The initial distribution of resistant isolates was based on surveillance data [3]. The major outcomes of interest were prevalence once equilibrium had been re-established and time to reach particular resistance thresholds in the population. We focused our threshold analyses on strains resistant to antibiotics B and C but not A and on strains resistant to all three antibiotics, since these are of clinical and public health importance. Time to reach 1% and 5% thresholds was calculated as time from model initiation to time at which strains displaying the resistance profile of interest represented n% or more of prevalent strains. The baseline comparator for all analyses was combination treatment of identified cases with antibiotics B and C. Simulations were run for 40 years, the point at which the system without a POC test had re-established equilibrium.

### Sensitivity analyses

We conducted sensitivity analyses for parameters describing the properties of the different resistant strains, test characteristics, and test coverage. We varied the fitness cost associated with resistance to antibiotic B (the antibiotic associated with the highest fitness cost for resistance) from 0 to 15%. We varied the relative fitness cost for antibiotics A and C from 0 to 1, and calculated the fitness cost for antibiotics A or C as:

> Fitness cost for antibiotic B x relative fitness cost for antibiotic A or C

 Additionally, we allowed the probability of *de novo* resistance acquisition to range from 10^−3^ to 10^−9^ per treatment event with each antibiotic.

In the main analysis we assumed that the POC test detected resistance with perfect sensitivity and specificity. We then varied test sensitivity and specificity to reflect that DNA-based tests may miss unrecognized mechanisms of resistance [16]. We assumed that the test properties for detecting resistance to each antibiotic were independent.

For the single resistance test, we assessed the impact of a fitness cost for antibiotic A resistance. For the 3-resistance POC test, we evaluated the alternate antibiotic selection strategy of treating with the antibiotic with the greatest barrier to resistance emergence (lowest probability of resistance emergence) when multiple treatment options were available.

## RESULTS

### Expected time course of resistance spread without a point-of-care test

Under our baseline assumptions of fitness costs and probabilities of *de novo* resistance emergence, as well as current patterns of resistance in the US population, the continued use of dual antibiotic treatment in the population was projected to result in >1% of isolates being resistant to both antibiotics within 12 years (Figure 2). With continued use of dual therapy in the population, this threshold surpassed 5% after an additional 3 years (year 15). Strains resistant to all three antibiotics took longer to become established, comprising >1% of all cases within 16 years. As resistant strains comprised a greater proportion of circulating strains and resulted in more treatment failures, the average time to successful treatment increased. Consequently, equilibrium prevalence was projected to increase approximately 3.5-fold in the modeled population, increasing from 2.1% at model start to 7.3% once the system re-equilibrated.

**Figure 2.**
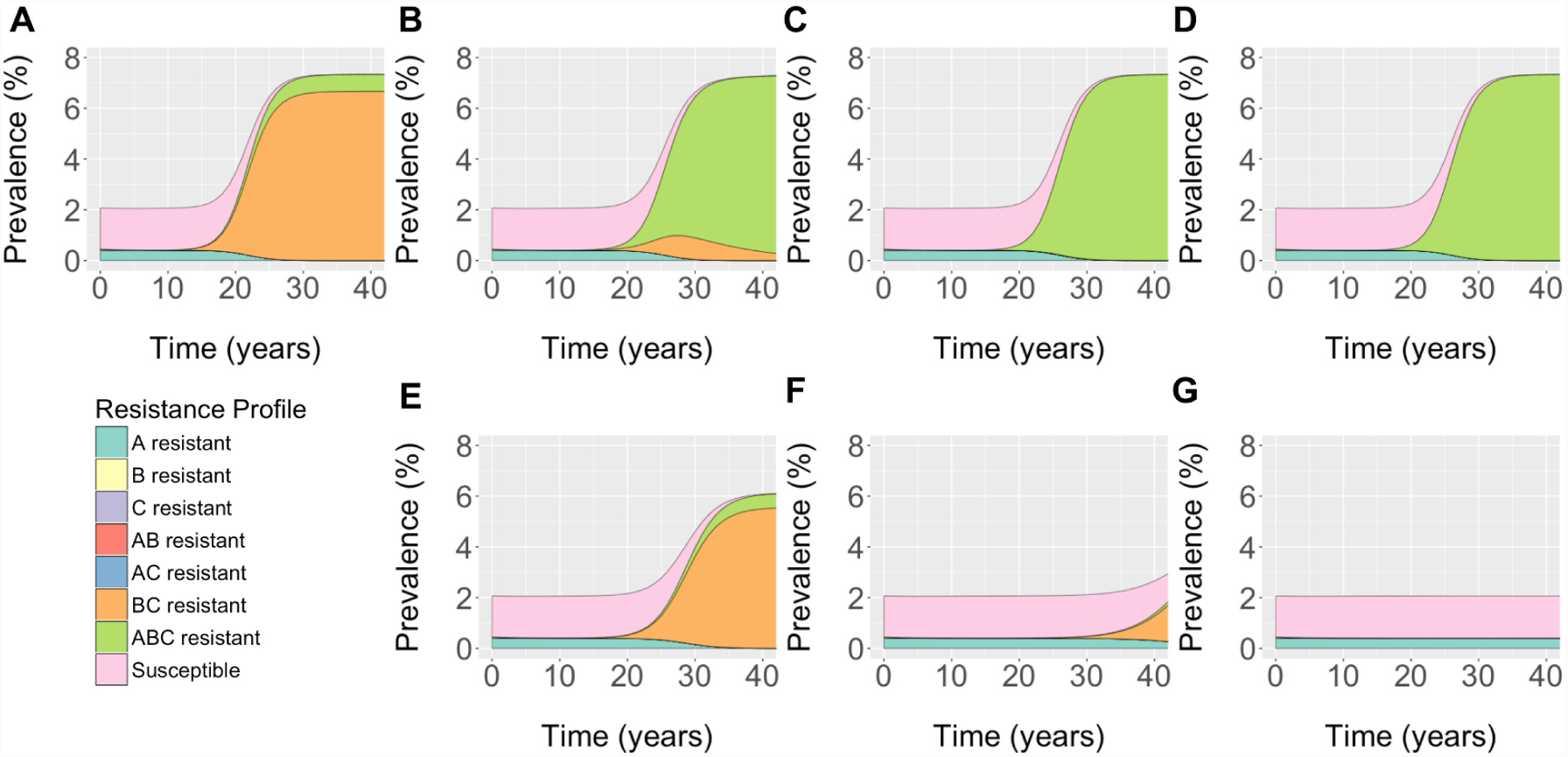
Projected impact of point-of-care tests on gonorrhea prevalence and resistance. Population prevalence and prevalence of different strains is shown in the face of (A) No POC testing, (B, E) 10%, (C, F) 25%, and (D, G) 50% of cases tested. Panels B-D show the results for a POC test for resistance to antibiotic A only. Panels E-G show result for a POC test for resistance to all three antibiotics. For the 3-resistance POC test, cases undergoing testing and displaying susceptibility to >1 antibiotic were treated with the antibiotic with the highest fitness cost associated with resistance acquisition. For both scenarios, all untested cases were treated in combination with antibiotics B and C. Results are shown for tests with perfect sensitivity and specificity.

### Impact of a POC test for resistance to a single antibiotic or three antibiotics

Under the baseline assumption of perfect test sensitivity and specificity, both POC test strategies had the identical probability that a BC resistant infection would be effectively treated (Figure S1). However, the single resistance POC test prompted a binary treatment decision based only on antibiotic A resistance status. Thus, all infections that tested resistant to A (including ABC resistant infections) were assigned BC treatment, making the probability of treating a triple resistant infection with an effective antibiotic zero, regardless of test coverage (Figure 3).

**Figure 3.**
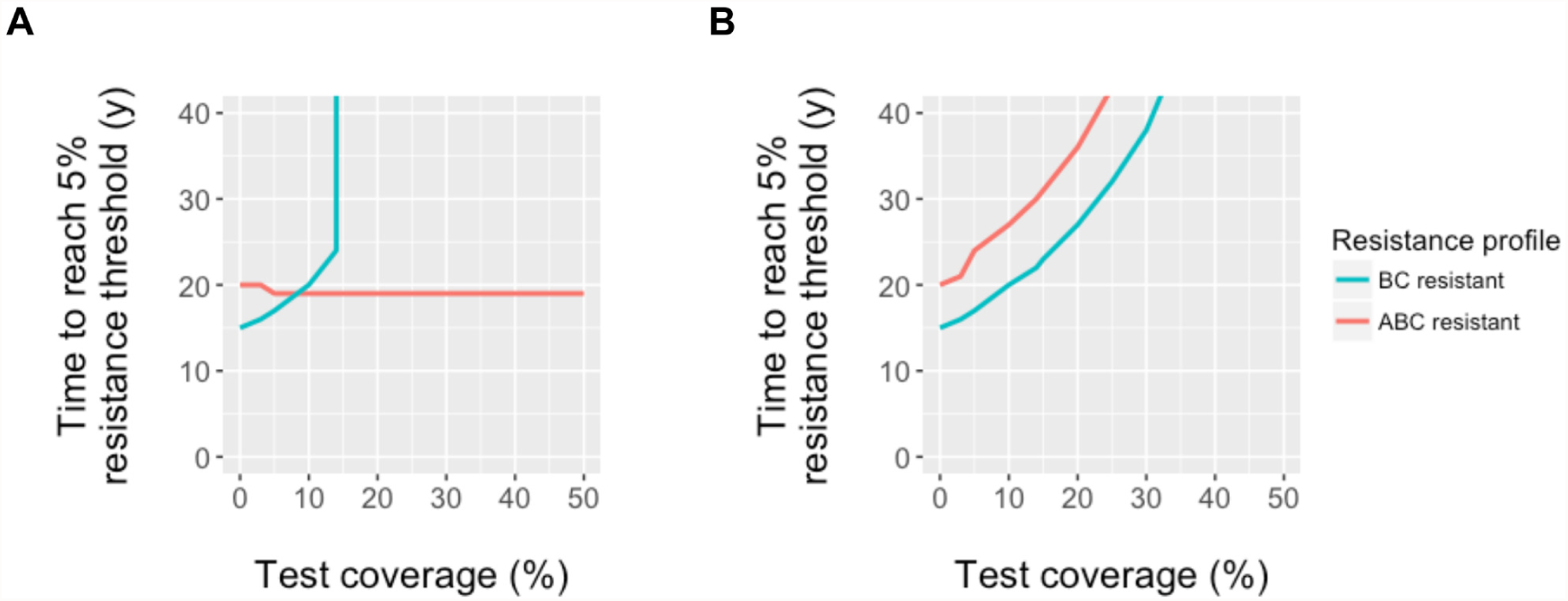
Time to resistance emergence with varying use of POC tests. Time for BC or ABC resistant strains to comprise 5% of prevalent gonorrhea isolates in the population with different point-of-care test use in the population. Results are shown for POC tests that identify resistance to (A) antibiotic A only or (B) all three antibiotics, with perfect sensitivity and specificity. Note that results are qualitatively similar for the 1% threshold, although times required to reach the threshold are reduced.

Unlike the base case, where BC resistance increased in the population, followed by ABC resistance, use of a single resistance POC delayed the spread of BC resistant gonococcal isolates but not ABC resistant isolates (Figure 2). Consequently, triple resistant strains could cross the resistance thresholds without BC resistance serving as a forewarning. For example, when test use was >20%, BC resistant isolates circulated in the population, but did not reach 1% of prevalent infections. Despite the reduction in BC resistance isolate transmission, the test did not reduce overall gonorrhea prevalence at equilibrium, relative to model projections in the absence of such a test, and had no or detrimental impact on triple resistance. With use of the test in 5% or more of cases, the time for ABC resistant strains to represent >5% of isolates was accelerated by 1 year, relative to no test use in the population.

By contrast, using a test that identifies resistance to all 3 antibiotics reduced overall equilibrium prevalence and delayed the spread of BC and ABC resistant strains in the population (Figure 2). With test use in >37% of identified cases, strains resistant to antibiotics B and C never reached 1% of prevalent cases within the 40-year time horizon. With higher test use, the rapid identification and treatment of BC and ABC resistant isolates allowed for isolates resistant to antibiotic A to persist. This reflected the assumption of no fitness cost associated with A resistance.

### Fitness cost associated with resistance to antibiotic A

Assuming a minor (1%) fitness cost for resistance to antibiotic A, triple resistant isolates were not projected to reach the 5% threshold during the 40-year time horizon when combination therapy was used to treat all identified infections. By contrast, use of the single resistance POC test (scenario I) resulted in this threshold being crossed in approximately 20 years once the test was used in greater than 5% of identified cases (Figure S2).

### Resistant strain properties

In sensitivity analyses, when fitness costs were relatively high (>15% for strains resistant to antibiotic B), resistant strains were outcompeted by other strains and did not reach the resistance thresholds, even without a POC test to guide antibiotic choice. When fitness costs were minor, the impact of the POC tests was diminished, in terms of the amount of time gained before resistance thresholds were crossed (Figure S3). As in our main analysis, the single resistance POC did not delay emergence of triple resistant isolates. Our results were less sensitive to fitness assumptions in scenarios with higher test coverage in the population (Figure S4 for triple resistance test, similar results for single resistance POC test).

Our findings were minimally sensitive to assumptions about the probability of resistance acquisition. Model projections changed only when probabilities were 10^−3^ per treatment event, larger values than would be considered biologically plausible [20]; even at this high level of resistance acquisition, the time to reach resistance thresholds was changed by 1-2 years compared to the base case estimates (Figure S5).

### Test characteristics

Test specificity only affected the projected impact of the single resistance test (Figure S6). For the triple resistance test, a false positive result would limit treatment options, but cases would still receive an effective treatment. For example, a BC resistant infection that was also falsely identified as A resistant, would receive an alternate treatment. In the case of the single resistance test, a BC resistant infection falsely identified as A resistant would be treated with BC, resulting in treatment failure.

Increased test sensitivity modestly increased the time until resistance thresholds were reached in the population (Figure S6). For example, when the test was used for 10% of cases, time to the reach the 5% BC resistance threshold increased by 4 years, as test sensitivity increased from 50 to 100%.

For the triple resistance test, relaxing the assumption that test sensitivity was identical for all three antibiotic resistant strains did not dramatically change our findings. As above, with reduced test sensitivities for detecting antibiotic A, B, and/or C resistance, the utility of the test for reducing gonorrhea burden and delaying the time until resistance thresholds were crossed was diminished (Figure S7).

### Alternate antibiotic selection approach with 3-resistance POC test

Basing antibiotic choice on probability of resistance acquisition on treatment rather than fitness costs associated with resistance did not have an impact on time to resistance emergence, regardless of assumed test sensitivity and test coverage, under the time horizon considered here (Figure S8).

## DISCUSSION

Using a mathematical model, we have shown that rapid diagnostics that report antibiotic susceptibility have the potential to extend the usefulness of existing antibiotics for treatment of gonorrhea as compared to the current guidelines for empiric two-drug treatment. Although most impactful when used in a large proportion of cases, even modest levels of use in the population can delay the establishment of resistance and reduce overall infection burden in the population.

Although our model projected a net benefit of a POC test, we found that a test for determining resistance to a single antimicrobial is not expected to delay, and may accelerate, emergence of triply resistant gonococcal isolates. The single antimicrobial test scenario was designed to replicate the potential effect of introduction of a rapid diagnostic for determining fluoroquinolone susceptibility. Although genomic and experimental analyses suggest there may not be a fitness cost associated with ciprofloxacin resistance [16, 21], our conclusions were unchanged even with a minor fitness cost. The failure of a single POC test to delay emergence of triply resistant isolates arises in part because all tested cases are treated appropriately except for triply-resistant infections, thereby reducing the burden of all other isolates and clearing the way for triply-resistant isolates. This finding underscores the importance of robust surveillance systems for understanding the landscape of resistance as such tests begin to be integrated into clinical care [10]. Surveillance data will be essential to detect and react to changes in resistance that may occur as a result of adoption of novel test technologies.

A POC test would enable clinicians to select antibiotic treatment from the set of drugs to which the pathogen is susceptible. Given that resistance to antibiotics can incur fitness costs, and that those costs differ by antibiotic, we evaluated a treatment strategy in which an infection with a strain susceptible to more than one antibiotic was treated with the antibiotic associated with the highest fitness cost, as inferred from population genomic and experimental data [16, 21]. We based this strategy on the reasoning that, should resistance to the antibiotic develop, a strain carrying a higher fitness cost would be less likely to persist in the population. Additionally, we evaluated the impact of guiding antibiotic selection by the probability of novel resistance emergence during the course of treatment. We note that both of these strategies delayed the emergence of resistance and rises in overall prevalence. Inferring the fitness costs of resistance, whether through population genomic data [16] or other means, would be an important parallel activity. Of note, the fitness costs need not be measured with precision. Instead, the relative ranking of costs per antibiotic guides selection.

Our study has several limitations. We used a deterministic model to determine how resistance would spread and assumed that the population was seeded with each of the resistant strains. Other modeling approaches that capture the stochastic nature of emergence and transmission are better suited to represent the inherent randomness in the emergence of resistance. However, use of a deterministic system allows us to draw initial inferences about the impact of POC test use, which can then be further explored using alternate modeling approaches. We did not explicitly model mixed infections (i.e., infections with multiple gonorrhea strains with different drug susceptibility profiles). However, imperfect test sensitivity in our model captures the impact of unrecognized resistant infections, whether they occur because a particular resistance marker is not included in the test, or because an individual has a mixed infection, with the resistant strain in low abundance. Actual treatment regimens used in the population are not 100% consistent with guidelines [3]. Similarly, resistance is not a binary trait [3]. More generally, as with all mathematical models, we made simplifying assumptions to describe gonorrhea transmission and resistance emergence and transmission. Given the early stages of POC test development, we sought to use this model to illustrate the potential impact of the use of such tests, and extensive sensitivity analyses demonstrate that the qualitative findings are robust under a range of assumptions and parameter values. We focused on a population experiencing a high burden of infection (MSM). As such, our analysis may be limited in generalizability to heterosexual populations, although trends in resistance in MSM often serve as harbingers of resistance in the broader population [11, 22]. The choice of an appropriate time horizon for this analysis was a compromise between allowing enough time for resistance to the current treatment regime to become established in the population, and acknowledging that novel treatment options may be introduced in the near future, limiting the policy relevance of long-term projections. Finally, our model did not consider changing fitness costs associated with antibiotic resistance, or declining test sensitivities over time, as might be expected if novel resistance mechanisms emerge. These represent complex questions and are an important avenue of exploration for future modeling studies.

Despite these limitations, this mathematical model demonstrates both the promise and potential need for caution associated with future POC tests for determining antibiotic susceptibility of gonococcal infections. The use of such tests cannot be done in isolation; continued real-time surveillance will be critical for guiding decision-making and monitoring resistance emergence.

## Acknowledgments

This project was funded by the U.S. Centers for Disease Control and Prevention, National Center for HIV, Viral Hepatitis, STD, and TB Prevention Epidemiologic and Economic Modeling Agreement (#5U38PS004642). The findings and conclusions in this report are those of the authors and do not necessarily represent the views of the Centers for Disease Control and Prevention.

## SUPPORTING INFORMATION

### Technical Appendix

**File S1.** Additional model details.

#### Supplementary Figures and Tables

**Figure S1.**
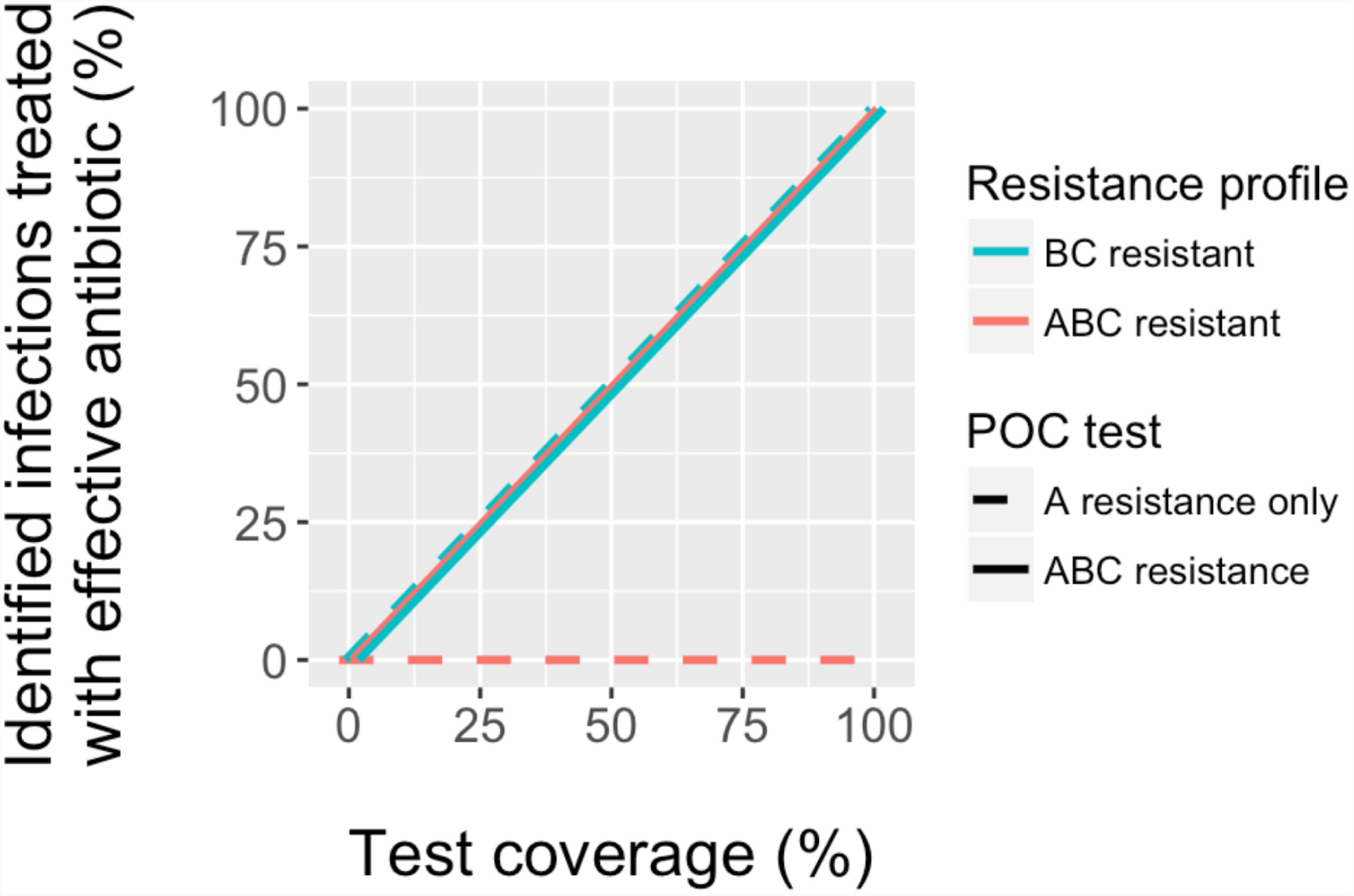
Probability of effective treatment with different POC tests. The probability that a BC or ABC resistant case receives an effective antibiotic is shown as test coverage is increased, with a POC test for A resistance only or resistance to all three antibiotics. Results are shown for tests with perfect sensitivity and specificity.

**Figure S2.**
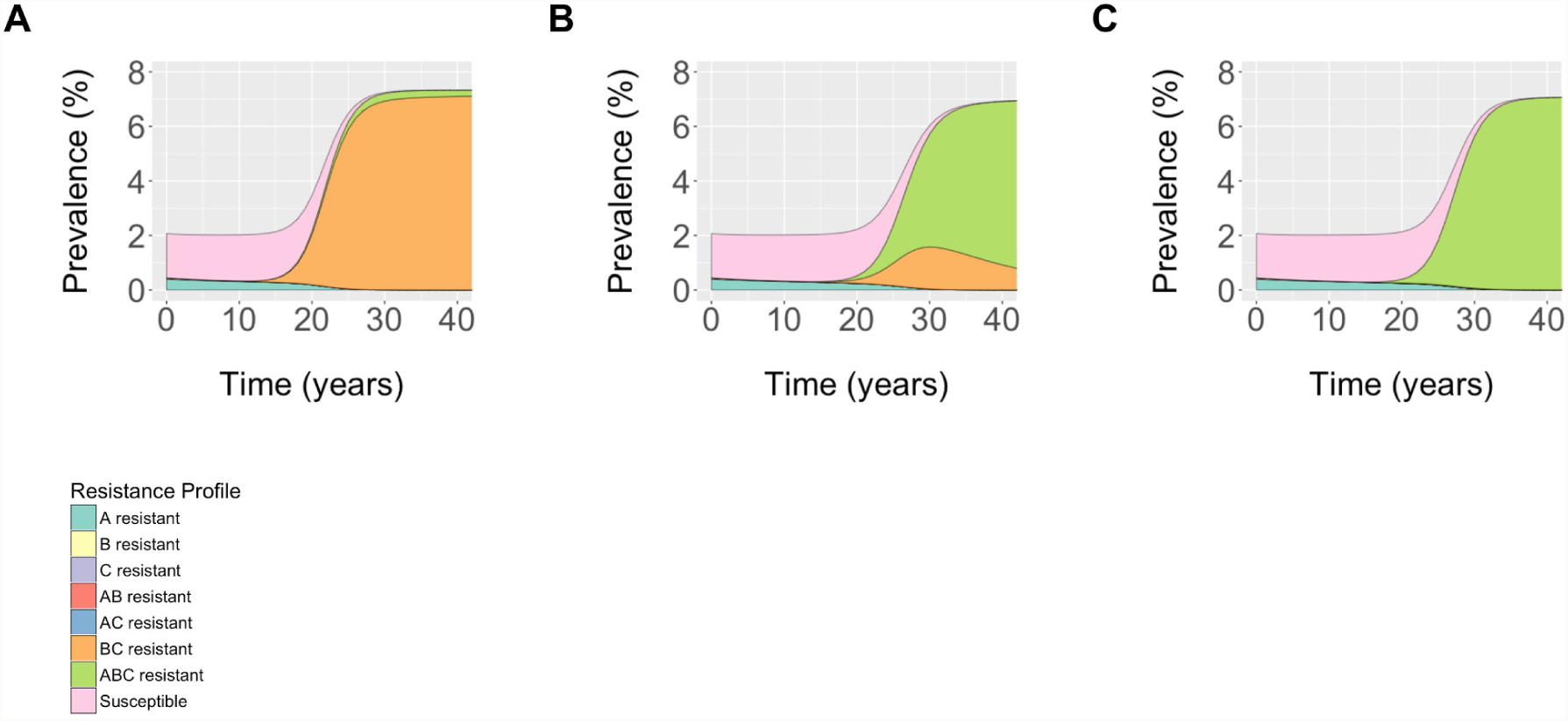
Model-projected gonorrhea prevalence with different levels of use of a point-of-care test for detecting single antibiotic resistance and reduced fitness for A resistant isolates. Population prevalence and prevalence of different strains is shown in the face of (A) No POC testing, (B) 10% of cases tested, and (C) 25% of cases tested. The fitness cost associated with antibiotic A resistance was 0.01. Fitness costs associated with resistance to the other antibiotics were unchanged from base case values. Cases undergoing testing and displaying susceptibility to antibiotic A were treated with antibiotic A; those displaying resistance to A were treated with combination BC. All untested cases were treated in combination with antibiotics B and C.

**Figure S3.**
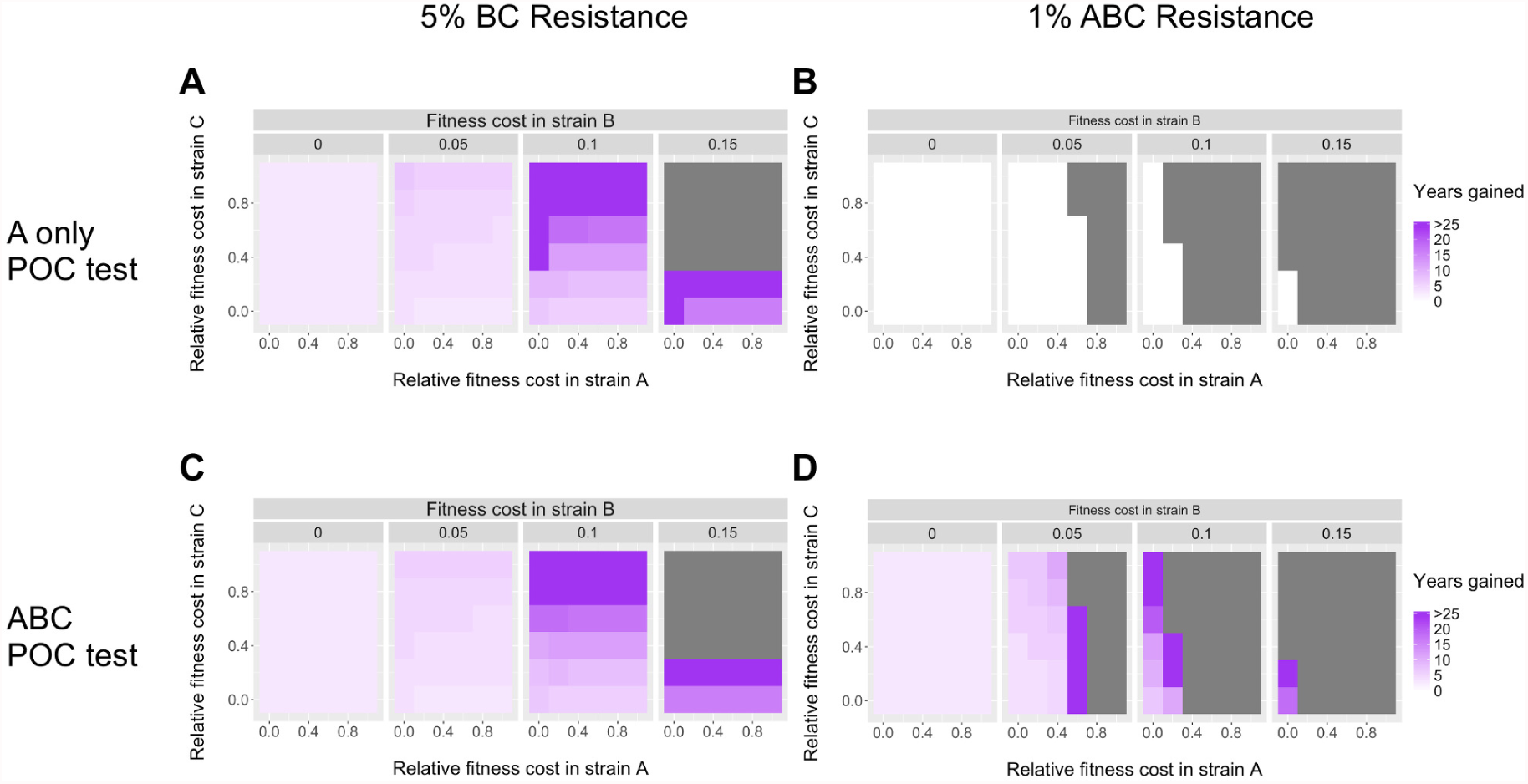
Model-projected gonorrhea prevalence with different levels of use of a point-of-care test for detecting single antibiotic resistance and reduced fitness for A resistant isolates. Impact of varying strain fitness on delays to resistance establishment with a point-of-care test. (A,C) Additional years for BC resistant strains to represent 5% of prevalent strains. (B,D) Additional years for ABC resistant strains to represent 1% of prevalent strains. A POC test determining resistance to (A,B) antibiotic A only, or (C,D) all three antibiotics was used for 10% of identified cases, with additional years gained calculated relative to no test use in the population. Detected cases are treated with combination BC therapy in the absence of a POC test. Fitness costs associated with resistance to antibiotic B are shown along the top panel. Fitness costs associated with resistance to antibiotics A and C are calculated as relative fitness costs, as described in Methods, with 0 representing no fitness cost associated with resistance, and 1 indicating an equivalent fitness cost as B resistance. Grey regions represent parameter combinations where the resistance threshold is not crossed within 50 years even in the absence of a test. Results are shown assuming perfect test sensitivity and specificity.

**Figure S4.**
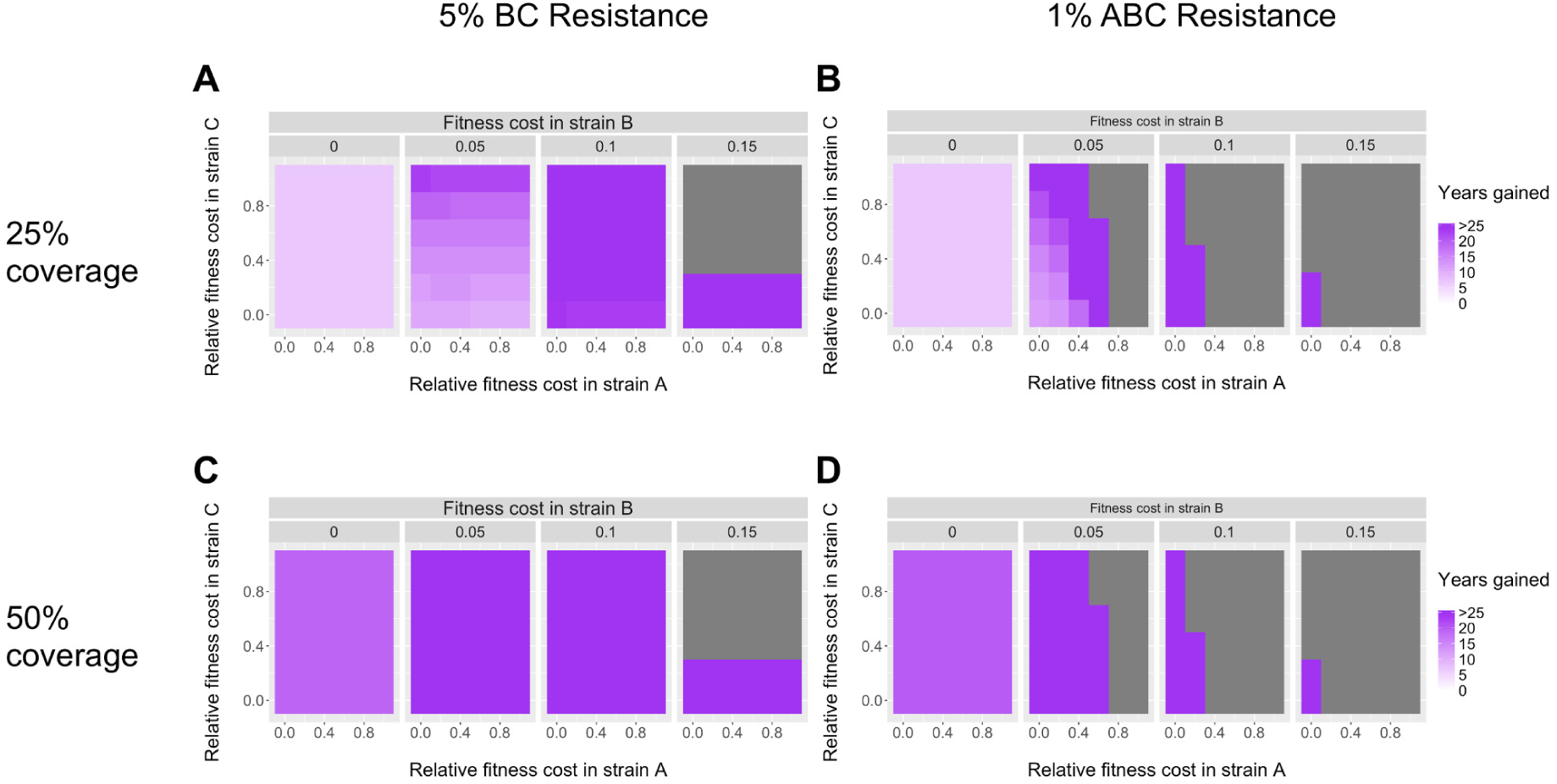
Impact of varying strain fitness on delays to resistance establishment with higher POC test coverage. Additional years for (A,C) BC resistant strains to represent 5% or (B,D) ABC resistant strains to represent 1% of prevalent strains, when a POC test determining resistance to all three antibiotics was used for (A,B) 25% or (C,D) 50% of identified cases. Additional years gained were calculated relative to no test use in the population. Detected cases are treated with combination BC therapy in the absence of a POC test. Fitness costs associated with resistance to antibiotic B are shown along the top panel. Fitness costs associated with resistance to antibiotics A and C are calculated as relative fitness costs, as described in Methods, with 0 representing no fitness cost associated with resistance, and 1 indicating an equivalent fitness cost as B resistance. Grey regions represent parameter combinations where the resistance threshold is not crossed within 50 years even in the absence of a test. Results are shown assuming perfect test sensitivity and specificity. Similar findings were observed for the single resistance POC test.

**Figure S5.**
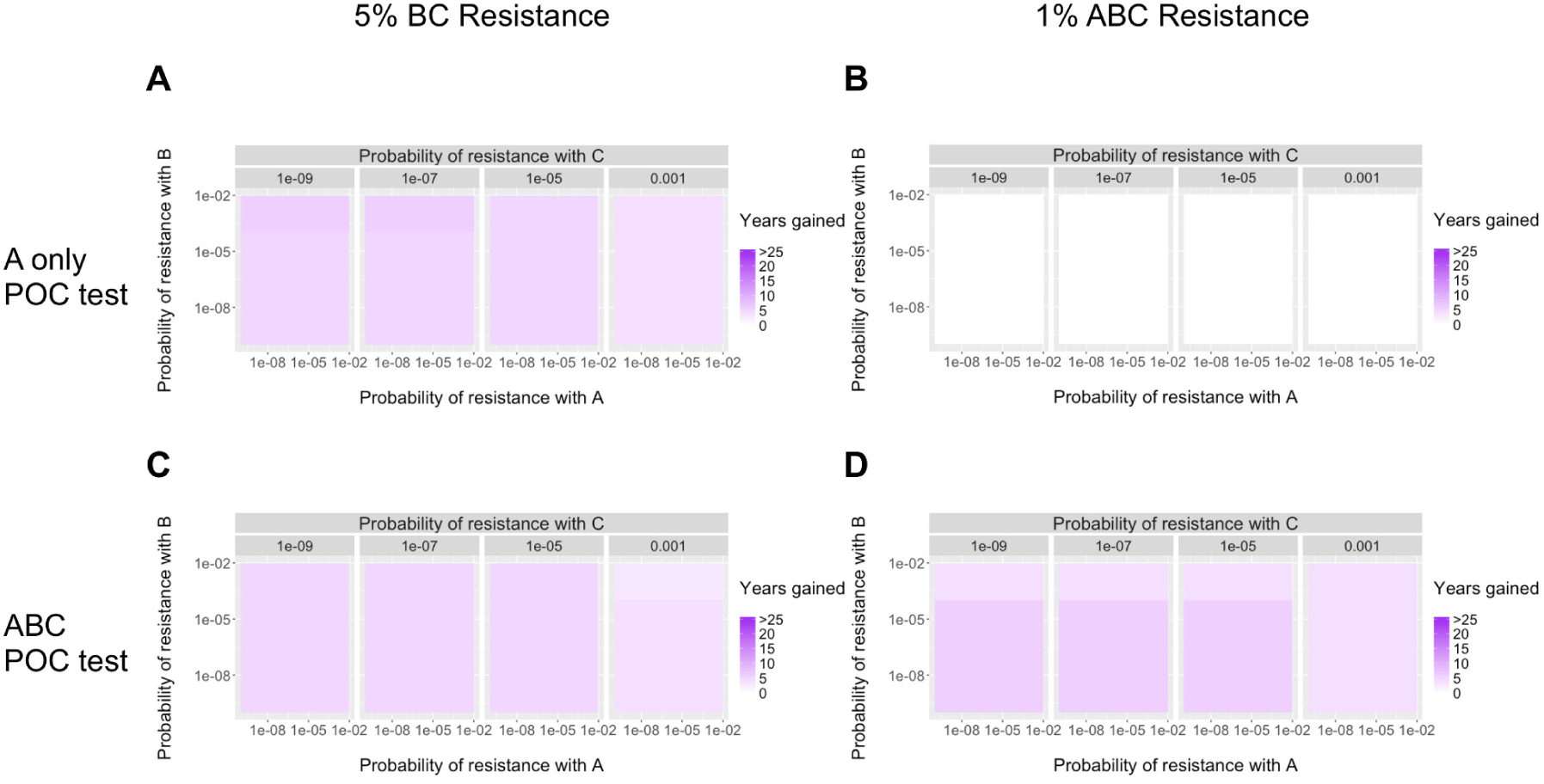
Impact of probability of resistance emergence on treatment on delays to resistance establishment with a point-of-care test. (A,C) Additional years for BC resistant strains to represent 5% of prevalent strains. (B,D) Additional years for ABC resistant strains to represent 1% of prevalent strains. A POC test determining resistance to (A,B) antibiotic A only, or (C,D) all three antibiotics was used for 10% of identified cases, with additional years gained calculated relative to no test use in the population. Detected cases are treated with combination BC therapy in the absence of a POC test. Probabilities of resistance emergence during treatment were varied for each antibiotic. Results are shown assuming perfect test sensitivity and specificity.

**Figure S6.**
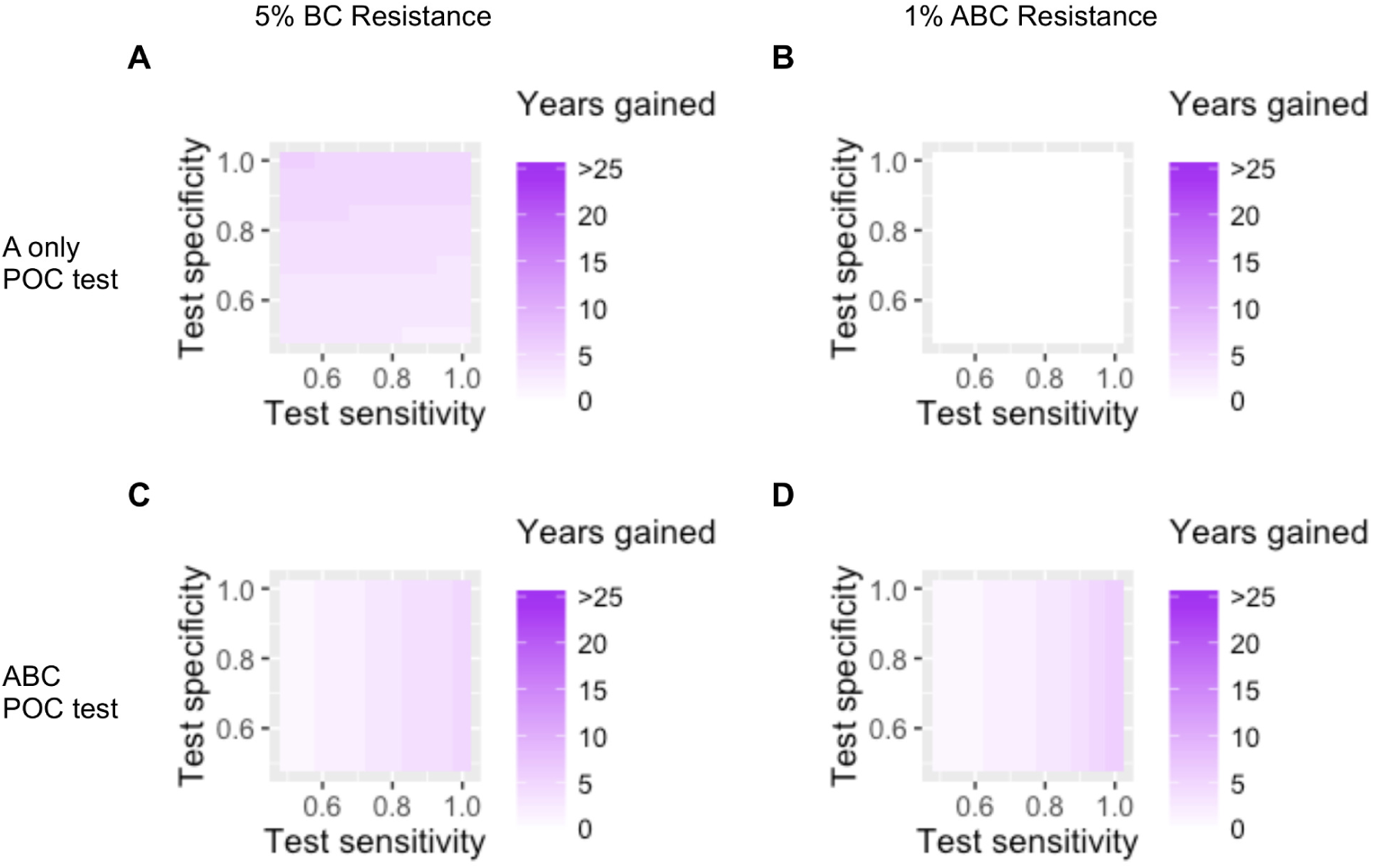
Impact of varying strain sensitivity and specificity on delays to resistance establishment with a point-of-care test. (A,C) Additional years for BC resistant strains to represent 5% of prevalent cases. (B,D) Additional years for ABC resistant strains to represent 1% of prevalent cases. A POC test determining resistance to (A,B) antibiotic A only, or (C,D) all three antibiotics was used for 10% of identified cases, with additional years gained calculated relative to no test use in the population. Detected cases are treated with combination BC therapy in the absence of a POC test. Sensitivity and specificity values represent test properties for detecting resistance to each antibiotic independently, as described in Methods.

**Figure S7.**
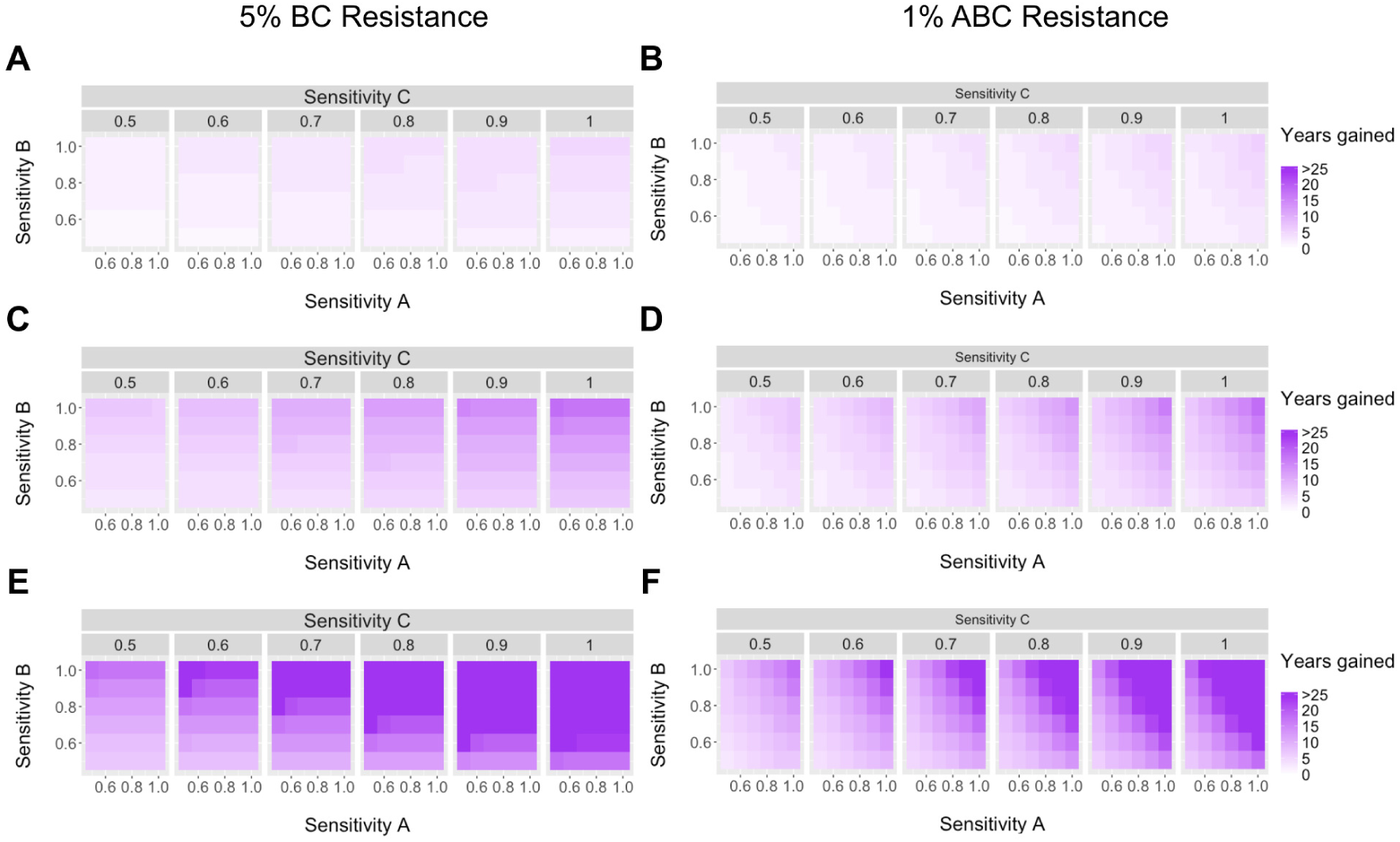
Impact of different test sensitivities for determining resistance to each of the three antibiotics included in the POC test. Test sensitivity for determining resistance to each antibiotic was varied from 50 to 100%. Additional years for (A,C, E) BC resistant strains to represent 5% or (B,D, F) ABC resistant strains to represent 1% of prevalent strains was calculated relative to no test use in the population. Results are presented assuming (A,B) 10%, (C,D) 25%, and (E,F) 50% test coverage. Detected cases are treated with combination BC therapy in the absence of a POC test.

**Figure S8.**
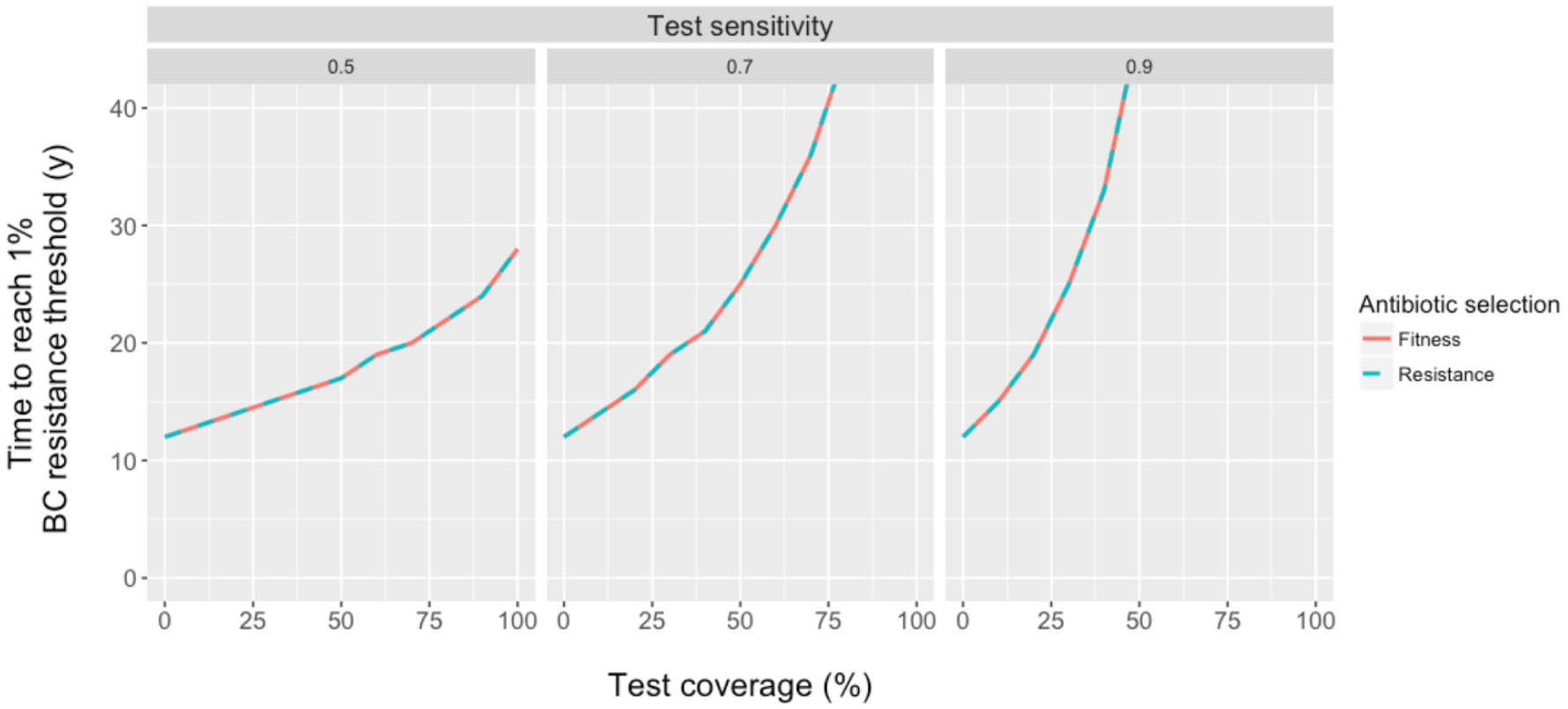
Effect of antibiotic selection strategy on POC test impact. When an infection was identified as susceptible to more than one antibiotic using the POC test for ABC resistance, treatment choice was based on either fitness cost (fitness) or probability of resistance acquisition on treatment (resistance). Results are shown for time until BC resistant isolates to represent greater than 1% of prevalent cases, with different test coverage and test sensitivity. Results are similar for time to ABC resistance.

**Table S1.**
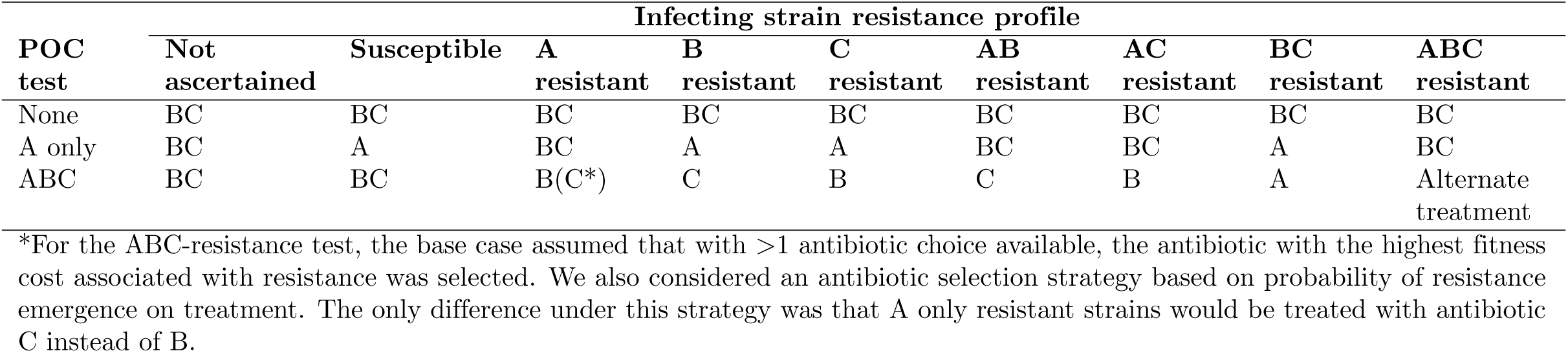
Antibiotic choice based on infecting strain resistance properties and testing strategy.

